# Xbra modulates the activity of linker region phosporlated Smad1 during *Xenopus* somitogenesis

**DOI:** 10.1101/2023.10.11.561863

**Authors:** Santosh Kumar, Zobia Umair, Ravi Shankar Goutam, Unjoo Lee, Jaebong Kim

## Abstract

The Bmp/Smad1 pathway plays a crucial role in developmental processes and tissue homeostasis. Mitogen-activated protein kinase (Mapk)/Erk mediated phosphorylation of Smad1 in the linker region leads to Smad1 degradation, cytoplasmic retention and inhibition of Bmp/Smad1 signaling. While Fgf/Erk pathway has been documented to inhibit Bmp/Smad1 signaling, several studies also suggests the cooperative interaction between these two pathways in different context. However, the precise role and molecular pathway of this collaborative interaction remain obscure. Here, we identified Xbra induced by Fgf/Erk signaling as a factor in a protective mechanism for Smad1. Xbra physically interacted with the linker region phosphorylated Smad1 to make Xbra/Smad1/Smad4 trimeric complex, leading to Smad1 nuclear localization and protecting it from ubiquitin-mediated proteasomal degradation. This interaction of Xbra/Smad1/Smad4 led to sustained nuclear localization of Smad1 and the upregulation of lateral mesoderm genes, while concurrently suppression of neural and blood forming genes. Taken together, the results suggests Xbra-dependent cooperative interplays between Fgf/Erk and Bmp/Smad1 signaling during lateral mesoderm specification in *Xenopus* embryos.

## Introduction

Mesoderm development is crucial event in vertebrate developmental biology mediating the formation of various tissues and organs such as muscles, skeletal structures, heart, blood, pronephros, and notochord (Kumano & Smith, 2002). In Xenopus embryos, mesodermal tissue patterning is regulated by signaling pathways like Wnt, TGF-β (nodal and Bmp), and Fgf (Amaya, Stein et al., 1993). The Fgf pathway is particularly vital, with Fgf2 being first identified Fgf as an inducer of mesoderm formation. Disruption of this pathway, using dominant negative Fgf receptors like XFD or Dnfr, leads to abnormal trunk and tail development(Amaya, Musci et al., 1991, Amaya et al., 1993, Kimelman & Kirschner, 1987). Subsequently, various Fgf ligands, including Fgf4 (referred to as embryonic Fgf or eFgf) and Fgf8 have been demonstrated for their ability to induce paraxial mesoderm in vivo (Fisher, Isaacs et al., 2002, Fletcher, Baker et al., 2006, Isaacs, Deconinck et al., 2007, Isaacs, Pownall et al., 1994, Isaacs, Tannahill et al., 1992, Standley, Zorn et al., 2001). Notably, in Xenopus embryo animal cap explants, eFgf has been noted as a potent inducer of mesoderm formation(Isaacs et al., 1994) and triggers the expression of the early pan-mesodermal marker Xbra (Amaya et al., 1993). Xbra, in turn, promotes mesoderm formation, and its mutual induction with eFgf forms an autocatalytic loop involving eFgf-Ras-Mapk-Xbra ensuring mesodermal fate maintenance and this induction process of Xbra is impeded in the absence of Fgf (Isaacs et al., 1994, Schulte-Merker & Smith, 1995, Yoon, Kim et al., 2014). Moreover, Fgf and Xbra expression in the blastula/gastrula marginal zone is essential for xmyoD expression and somite formation (Harvey, 1992, Hopwood, Pluck et al., 1989) disruption of the Fgf pathway results in the loss of Xbra, hindering somite formation and expanding the blood island. (Kumano & Smith, 2002).

Bone Morphogenetic Protein 4 (Bmp4), part of the transforming growth factor beta (TGFβ) superfamily, plays a crucial role in mesoderm formation and specification (Hoppler & Moon, 1998). Bmp4 signals through the Smad1/5 pathway, where Bmp receptor (BmpR) phosphorylates Smad1/5 c-terminal promoting interaction with co-Smad Smad4. This complex then moves to the nucleus, regulating various mesodermal and epidermal genes (Chen, Zhao et al., 2004, Shi & Massague, 2003). Research, including gain and loss-of-function studies, highlights Bmp4 critical role in somite formation and tailbud outgrowth in frog and chick embryos (Beck, Whitman et al., 2001, Sharma, Shafer et al., 2017). Disruption of Bmp activity in zebrafish is linked to abnormal ventral fin and tail mesoderm development (Pyati, Webb et al., 2005, Stickney, Imai et al., 2007). In mice, Bmp signaling is crucial for maintaining mesoderm progenitor homeostasis in the tailbud, activating key factors in the mesoderm molecular network (Winnier, Blessing et al., 1995). In Xenopus embryos, evidence suggests that Bmp4 concentration influences dorsal-ventral mesodermal structures. Low Bmp mRNA concentrations upregulate xmyf5, xvent1, and xvent3 expression dorsally, while high concentrations inhibit xmyf5 and induce xvent1 and xvent3 throughout the marginal zone. Low doses of Dnfr (receptor antagonist) promote xmyf5 expression and suppress xvent1 in the ventral marginal zone, while high doses lead to the loss of xmyf5 and xvent3 expression, upregulating dorsal genes throughout the marginal zone (Dale & Jones, 1999) indicating need of fine balanced amount of Bmp4 for diverse mesoderm tissue patterning.

Mesodermal specification mediated by Bmp signaling requires input from additional pathways including the Fgf/Mapk/Xbra cascade. For instance, the Bmp target gene xvent1 showed increased expression in response to eFgf, leading to the absence of blood island formation (Xu, Ault et al., 1999). Fgf’s cooperative role has also been observed in limb and kidney development (Godin, Robertson et al., 1999, Makanae, Mitogawa et al., 2014). In our prior research, we found that Smad1 and Xbra collaborate to enhance xvent1 expression through the formation of a Smad1/Xbra complex binding to the xvent1 promoter (Kumar, Umair et al., 2018, Yoon et al., 2014). Conversely, Fgf input via Mapk at the Smad1 level has been found to be inhibitory, particularly in neural specification (De Robertis & Kuroda, 2004). Neural inducers, such as Fgf8 and IGF phosphorylate the linker region of Smad1 at four conserved serine sites by activating tyrosine kinase receptors. This phosphorylation event inhibits Smad1 nuclear localization while promoting neural induction (De Robertis & Kuroda, 2004, Pera, Ikeda et al., 2003). Furthermore, Fgf-mediated linker region phosphorylation primes Smad1 for GSK3β-mediated phosphorylation, leading to smurf1 and E3-mediated polyubiquitination and degradation or cytoplasmic retention (Sapkota, Alarcon et al., 2007). These distinct set of findings highlight the contextual cross-talk between Fgf/Mapk/Xbra and Bmp4/Smad1 pathways during neural and mesodermal specification. While the Fgf-Mapk cascade’s input to Smad1 in neural differentiation is well-studied, the cooperative relationship between Fgf-Mapk-Xbra and Bmp/Smad1 during mesoderm specification is intriguing and require further exploration. The precise function and underlying molecular mechanisms of the Xbra-Smad1 cross-talk remain to be comprehensively addressed.

Here we identified Xbra, generated by eFGF/Mapk signaling, plays a crucial role in regulating Smad1 activity and stability in the presence of Bmp4 and Mapk/Erk-mediated phosphorylation input. We found Xbra as an essential factor required for the nuclear localization and transcriptional activity of Smad1 which is phosphorylated at both the c-terminal (by Bmp receptors) and linker region (by Fgf/Mapk/Erk). We also found that in the presence of Xbra linker region phosphorylated Smad1 is protected from polyubiquitination leading to enhanced stability over time, even in the presence of exogenous GSK3β. This coexistence of eFgf/Mapk/Erk and Bmp4 input at Smad1 in the presence of Xbra induced the expression of various mesodermal specifiers including *xvent1*, xvent3, *myoD*, *myf5*, *muscle-actin*, and *myogenin*. Additionally, our findings highlighted the importance of Smad4 in forming the Xbra/Smad1 complex, facilitating its nuclear localization and subsequent transcriptional activity. Overall, our results shed light on the intricate interplay between eFgf/Mapk/Erk/Xbra and Bmp/Smad1 in lateral mesoderm specification.

## Results

### eFgf enhances Smad1 nuclear localization and lateral mesoderm specifier expression in Bmp4 environment

In this study, we aimed to investigate the influence of eFgf on Smad1 nuclear localization and its impact on the expression of lateral mesoderm specifiers in the presence of Bmp4. We initially examined the nuclear localization of endogenous Smad1 in animal cap explants. In the explants from embryo injected with Bmp4 and eFgf, we observed a significant increase in Smad1 nuclear localization compared to injected with Bmp4 alone (Fig. 1A). To confirm these findings, we repeated this experiment using flag-tagged smad1 (flag-smad1), which also showed nuclear localization in the presence of Bmp4 and eFgf (Fig. 1B). To assess the impact of eFgf and Bmp4 on gene expression within the animal cap explants, we conducted qPCR assays for a range of early and late genetic markers. In the presence of both Bmp4 and eFgf, we observed a significant upregulation of early somite markers, including *xbra*, *myf5*, and *myoD* (Fig. 1 C, D and E). Additionally, *xvent1* and *xvent3* were induced by eFgf and Bmp4 (Fig. 1 F and G). However, the co-injection of eFgf and Bmp4 did not lead to any significant increase in the expression of early neural markers like *sox3* and *otx2* (Fig. 1 H and I). Notably, the early ventral mesodermal marker *gata2* was also not upregulated in this context (Fig. 1 J). Furthermore, evaluating the late mesodermal markers the co-injection of eFgf with Bmp4 resulted in a significant elevation in the expression of skeletal muscle markers, *myogenin*, and *muscle-actin* (*m-actin*) Fig. 1 K and L). However, expression of cardiac muscle markers *myocardin* and *cardiac-α-actin* remained unaltered or downregulated (Fig. 1 M and N). Neural marker *n-cam* also exhibited no significant change in the presence of eFgf and Bmp4 (Fig. 1 P). The ventral blood island marker *globin* showed diminished expression in the presence of eFgf and Bmp4, while Bmp4 alone notably elevated the expression of *gata2* and *globin* (Fig.1 O and J). In summary, our results indicate that eFgf enhances Bmp/Smad1 activity by promoting the nuclear localization of Smad1. This cooperative effect of eFgf and Bmp4 leads to the induction of myogenic markers in the animal cap explants, while sparing early neural and ventral blood markers.

**Fig. 1.**
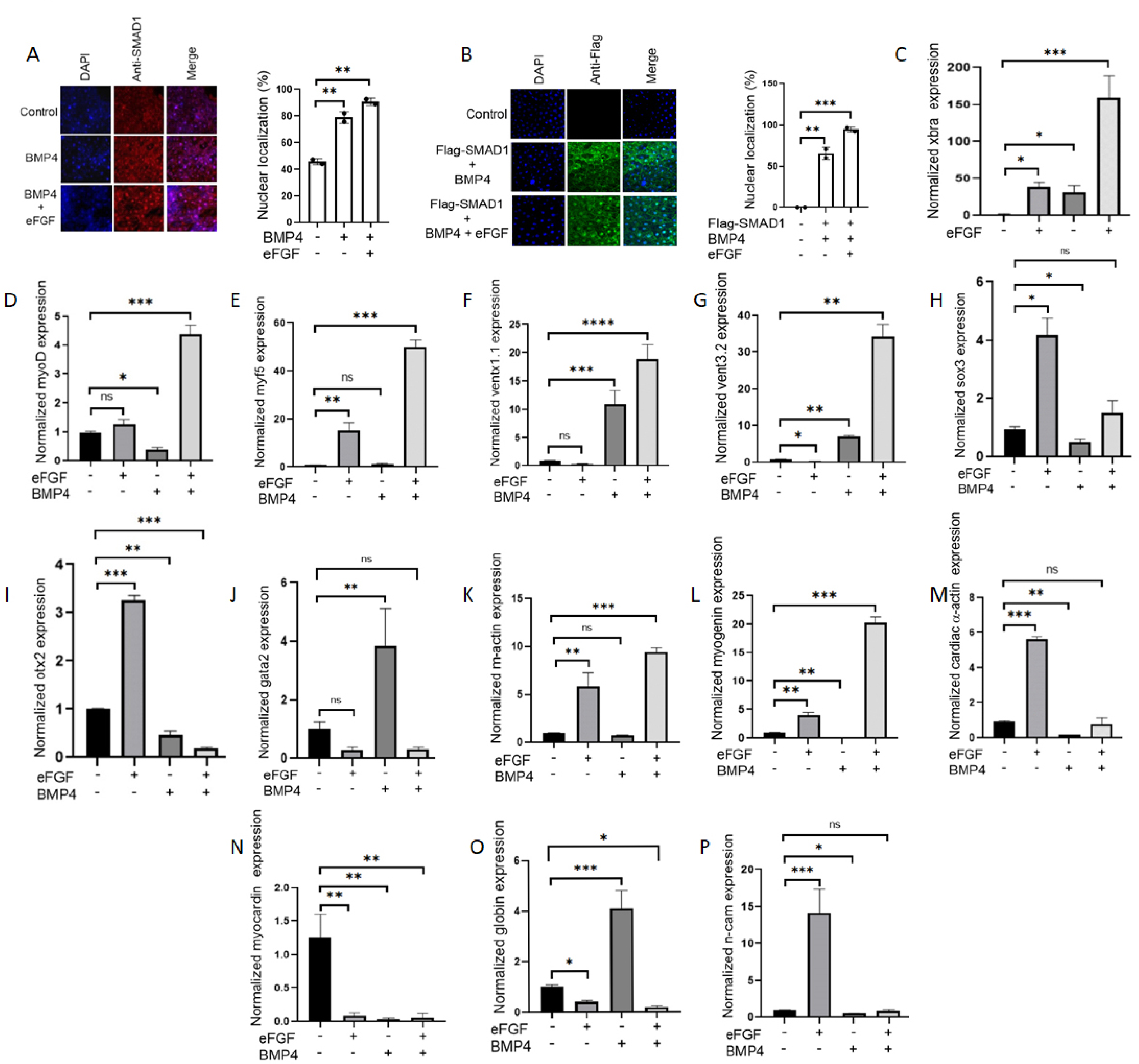
Influence of eFGF on smad1 nuclear localization and gene expression in BMP4-treated animal cap explants. (A) Subcellular localization of smad1 in Xenopus animal cap explant in the presence of control, BMP4 and BMP4 with eFGF injected. Animal cap dissected at 9.5 stage and grown until stage 10.5, fixed and then subjected to immunofluorescence assay with primary antibody against endogenous smad1 and anti-rabbit Alexa-fluor 594 secondary antibody (Invitrogen) was used. Blue: DAPI, Red: endogenous smad. Scale bar 20X. This experiment was repeated two times with consistent results. Right: Statistic data of smad1 nuclear translocation ratio. (B) Subcellular localization of flag-smad1 in Xenopus animal cap explant in the presence of control flag-smad1, flag-smad1 with BMP4 and flag-smad1 with BMP4 and eFGF injected. Animal cap dissected at 9.5 stage and grown until stage 10.5, fixed and then subjected to immunofluorescence assay with primary antibody against endogenous smad1 and anti-mouse Alexa-fluor 488 secondary antibody (Invitrogen) was used. Blue: DAPI, Green: flag tag. Scale bar 20X. This experiment was repeated two times with consistent results. Right: Statistic data of Flag-smad1 nuclear translocation ratio. (C-J) qRT-PCR Analysis of the gene expression levels of early markers in animal cap explants from embryos injected with eFGF and BMP4 together or separately. qRT-PCR values were determined from the ΔΔCt for the target genes relative to odc. Significantly different results for treatments used one-way ANOVA with Graph Pad Prism; P < 0.05. (K-P) qRT-PCR Analysis of the gene expression levels of late markers in animal cap explants from embryos injected with eFGF and BMP4 together or separately. qRT-PCR values were determined from the ΔΔCt for the target genes relative to odc. Significantly different results for treatments used one-way ANOVA with Graph Pad Prism; P < 0.05.

### eFgf and Bmp4 input to smad1 synergistically enhance Smad1/Xbra complex formation

Previous researches has established that Fgf, including eFgf, induces Xbra formation through the Fgf/Mapk/Xbra autocatalytic loop and promotes *xvent1* expression while suppressing blood formation via Xbra-Smad1 complex formation (Amaya et al., 1993, Kumar et al., 2018, Xu et al., 1999, Yoon et al., 2014). It has also been reported that Fgf, via Mapk/Erk, phosphorylates the linker region of Smad1 (Pera et al., 2003) . In this study (Fig. 1), we found the cooperative effects of eFgf and Bmp and their influence on Smad1 nuclear localization and the upregulation of skeletal muscle markers. To assess whether eFgf promote Mapk/Erk-mediated phosphorylation to Smad1 linker region and does this phosphorylation play any role in Smad1-Xbra complex formation, we conducted immunoprecipitation experiments in the presence and absence of Fgf/Mapk input. As depicted in Fig. 2A, we observed that the presence of eFgf significantly promoted complex formation between endogenous Smad1 and myc-Xbra. However, this complex formation was notably reduced in the presence of the Mapk/Erk inhibitor, U0126. Similarly, as shown in Fig. 2B, the complex formation between flag-Smad1 and myc-Xbra was also reduced in the presence of U0126, indicating the role of Fgf/Mapk signaling in promoting this interaction. Previous knowledge suggested that Fgf/Mapk phosphorylates Smad1 at specific sites in the linker region (S187, S196, S206, and S214) (Chen & Wang, 2009). To confirm if eFgf has a similar phosphorylation effect on Smad1, we conducted Western blot analysis which demonstrates that the injection of eFgf significantly increased the phosphorylation of Smad1 at LS206 and L214 site (Fig. 2 C). To further investigate the role of linker region phosphorylation of Smad1 in promoting Smad1-xbra complex formation, we constructed two mutant forms of flag-smad1. The first mutant, flag-Smad1(4SD3SD), was designed with constitutive phosphorylation at the SSxS site at the C-terminus (BmpR target site) and the four Fgf/Erk-targeted serine phosphorylation sites in the linker region changed to DDDD (constitutively phosphorylated form). The second mutant, flag-Smad1(4SA3SD), had the SSxS site at the C-terminus (BmpR target site) changed to DDxD (constitutively phosphorylated form), while the four Fgf/Erk-targeted serine phosphorylation sites (SSSS) in the linker region were mutated to AAAA (Fig. 2 D). As anticipated, our results showed higher complex formation between flag-Smad1 (4SD3SD) and myc-Xbra compared to the wild-type flag-Smad1, whereas complex formation between flag-Smad1(4SA3SD) and myc-Xbra was significantly reduced (Fig. 2E). These findings strongly suggest that Mapk/Erk-mediated phosphorylation of the Smad1 linker region plays supportive role in inducing Xbra-Smad1 complex formation. In summary, our study demonstrates that eFgf and Bmp cooperation leads to Smad1 nuclear localization and the upregulation of skeletal muscle markers. Additionally, we provide evidence that Mapk/Erk-mediated phosphorylation at specific sites in the Smad1 linker region is required for Smad1-Xbra complex formation.

**Fig. 2.**
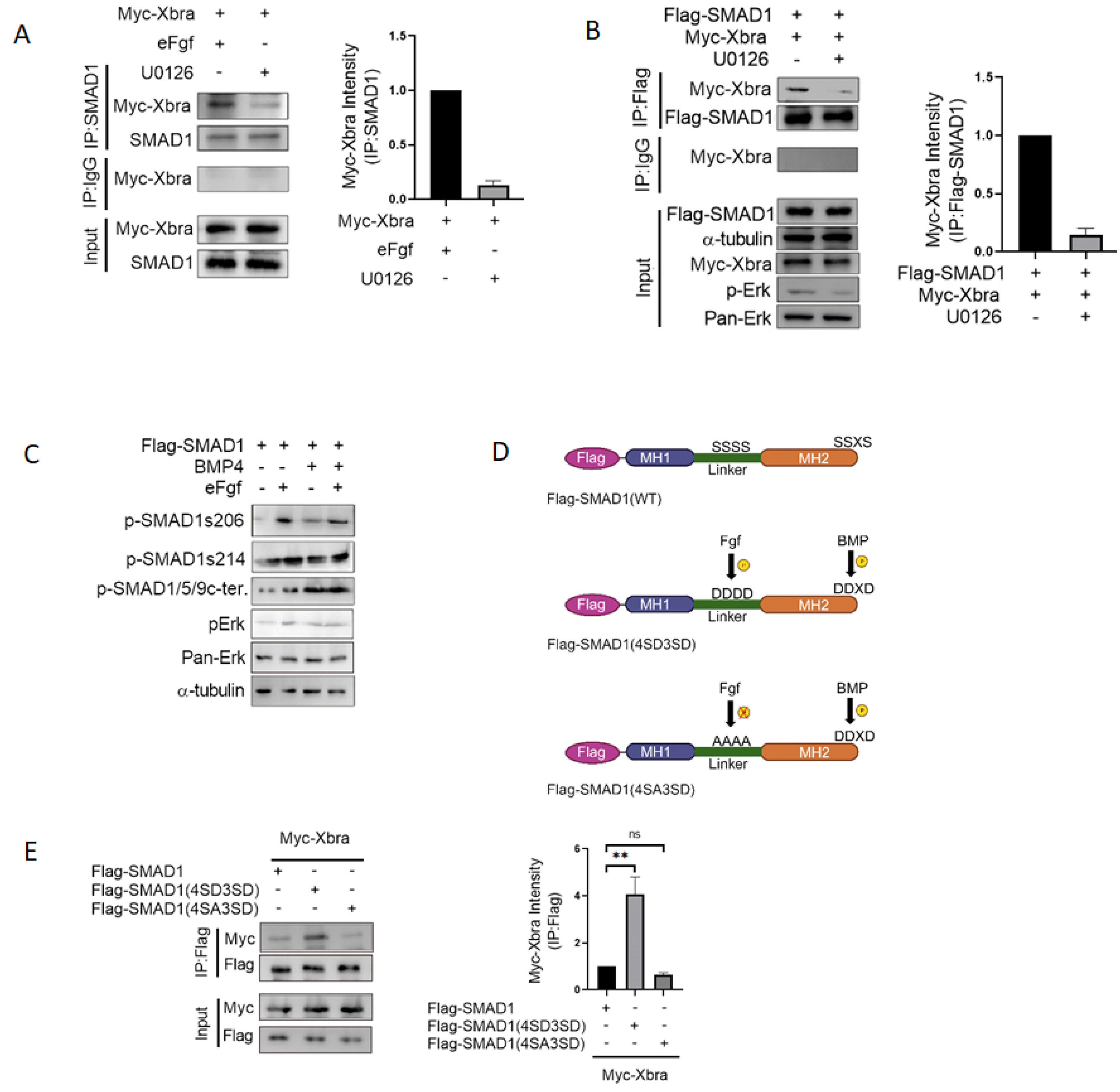
Synergistic enhancement of Smad1/Xbra complex formation through eFgf and Bmp4 stimulation. (A) Animal cap from embryos injected with myc-xbra and eFGF either treated with U0126 or control DMSO and harvested for co-immunoprecipitation analysis using endogenous smad1 and anti-myc antibodies. Right: Myc-Xbra level were quatified by photo shop software. This experiment was repeted two times with consistemce results. (B) Animal cap from embryos injected with flag-smad1 and myc-xbra either treated with U0126 or control DMSO and harvested for co-immunoprecipitation analysis using anti-flag and anti-myc antibodies. α-tubulin served as loading control. . Right: Flag-smad1 level were quatified by photo shop software. This experiment was repeted two times with consistemce results. (C) Immunoblotting analysis of total lysates from animal cap explant from embryo injected with flag-smad1 together with or without eFGF and BMP4 lysates were analyzed by immunoblotting using p-smad1linker206, p-smad1linker214, p-smad1/5/9c-terminal and p-Erk antibody. Pan Erk and α-tubulin served as a loading control. (D) Smad1 phospho-mutant forms, constitutively active FGF- and BMPR-mediated phosphorylated form of flag-smad1(4SD3SD), and constitutively inactive FGF- and active BMPR-mediated phosphorylated form of flag-smad1(4SA3SD) along with wild type. (E) Embryos injected with flag-smad1 (WT), flag-smad1(4SD3SD) or flag-smad1(4SA3SD) with myc-xbra and harvested for co-immunoprecipitation analysis using anti-flag and anti-myc antibodies. Right: Myc-Xbra level were quatified by photo shop software. This experiment was repeted two times with consistemce results.

### Fgf and Bmp mediated phospho-mimetic smad1 mutant require Xbra for nuclear localization and induction of lateral mesoderm markers

To further support our findings, we next investigated the nuclear localization and transcriptional activity of the constitutively BmpR and eFgf mediated phosphorylated mutant form of Smad1 (flag-smad1(4SD3SD)) in the presence of Xbra. To elucidate these, we injected flag-smad1(4SD3SD) both with and without myc-Xbra and performed immunofluorescence analysis using animal cap explants. As depicted in Figure 3A, flag-smad1(4SD3SD) exhibited negligible nuclear localization when administered alone, but in the presence of Xbra, a significant portion was observed within the nucleus. To further evaluate the transcriptional activity of flag-smad1(4SD3SD) in the presence of Xbra, we conducted qPCR assays with animal cap explants. In presence of Xbra and flag-smad1(4SD3SD) condition, we observed a substantial upregulation in the expression of myogenic markers, including *xvent1*, *xbra*, *myoD*, and *myf5* at the early developmental stage, and *myogenin* and muscle-actin (m-actin) at the later stage. Whereas, flag-smad1(4SD3SD) alone failed to induce any of these markers (Fig. 3B-G). Additionally, neither alone flag-smad1(4SD3SD) nor with Xbra could induce any cardiac muscle markers, such as *cardiac-α-actin* and *myocardin* (Fig. 3H, I). Furthermore, *globin* expression was also not induced by flag-smad1(4SD3SD) in the presence or absence of Xbra; it was upregulated only in the presence of WT flag-smad1 (Fig. 3J). Of note, flag-smad1(4SD3SD) alone exhibited *n-cam* expression and the presence of Xbra attenuated this induction (Fig. 3K). This expression of the neural marker *n-cam* in the overexpressed flag-smad1(4SD3SD) condition may be attributed by the inhibition of Bmp signaling output. In summary, our findings indicate that the dual input of eFgf/Mapk and Xbra to Smad1by Fgf pathway through phosphorylation of the Smad1 linker region and the formation of a Smad1-Xbra complex, leads to the nuclear localization of the Smad1-Xbra complex and its subsequent transcriptional activation, resulting in the induction of myogenic markers.

**Fig. 3.**
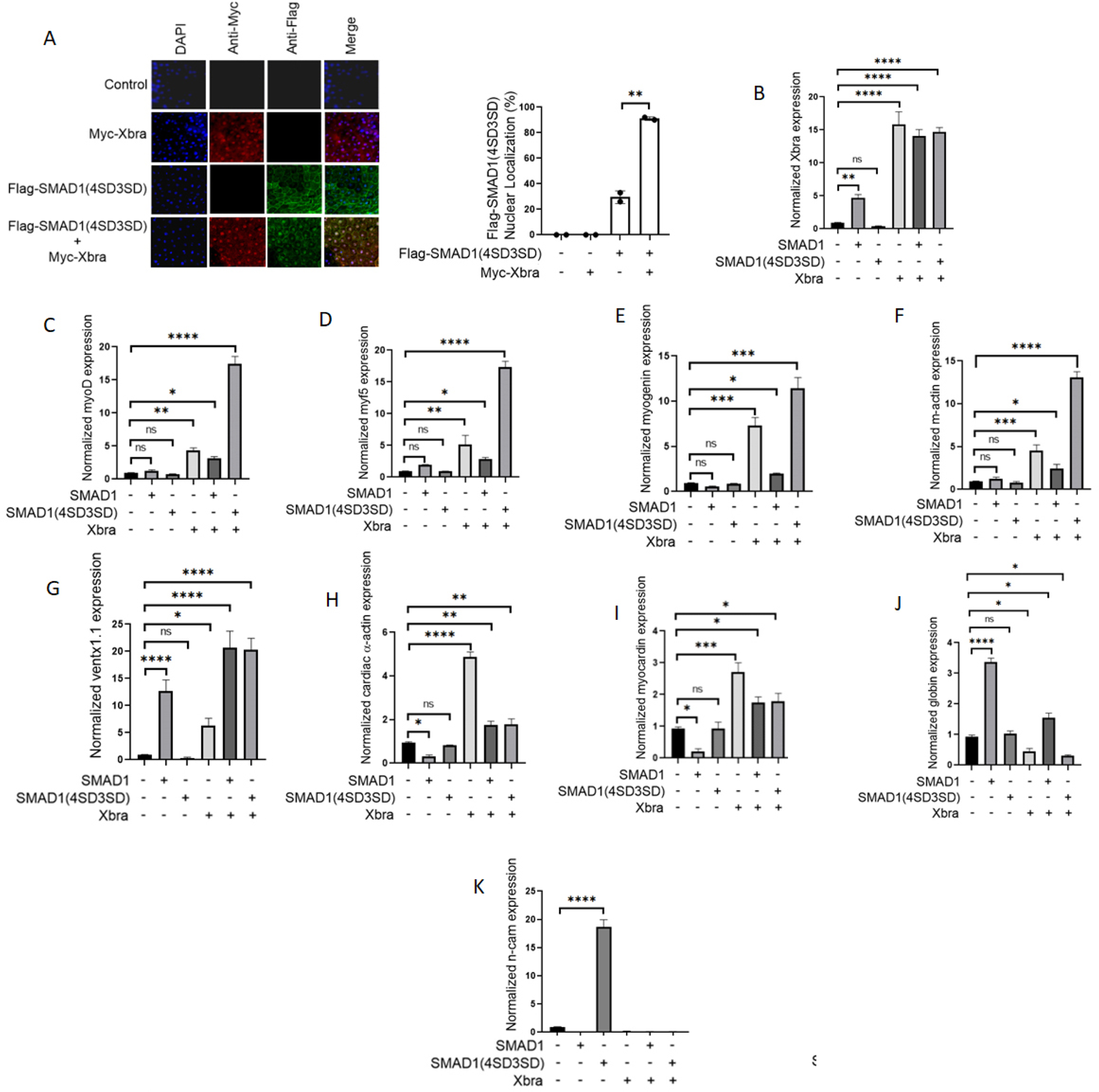
The nuclear localization and induction of lateral mesoderm markers by the phospho-mimetic Smad1 mutant, mediated by Fgf and Bmp, require Xbra. (A) Subcellular localization of flag-smad1(4SD3SD) in Xenopus animal cap explant from embryos injected with flag-smad1(4SD3SD) and myc-xbra together and separately. Animal cap dissected at 9.5 stage and grown until stage 10.5, fixed and then subjected to immunofluorescence assay using primary antibody against flag-tag and myc-tag. Anti-mouse Alexa-fluor 488 and anti-rabbit Alexa fluor 594 secondary antibody (Invitrogen) was used. Blue: DAPI, Green: flag tag, Red: myc-tag. Scale bar 20X. This experiment was repeated two times with consistent results. Right: Statistic data of smad1 nuclear translocation ratio. (B-E) qRT-PCR Analysis of the gene expression levels of early markers in animal cap explants from embryos injected with smad1(WT) and smad1(4SD3SD) with or without xbra. qRT-PCR values were determined from the ΔΔCt for the target genes relative to odc. Significantly different results for treatments used one-way ANOVA with Graph Pad Prism; P < 0.05. (F-K) qRT-PCR Analysis of the gene expression levels of late markers in animal cap explants from embryos injected with smad1(WT) and smad1(4SD3SD) with or without xbra. qRT-PCR values were determined from the ΔΔCt for the target genes relative to odc. Significantly different results for treatments used one-way ANOVA with Graph Pad Prism; P < 0.05.

### Xbra functions to stabilize endogenous smad1 and a phospho-mimetic smad1 mutant by inhibiting its polyubiquitination and subsequent degradation

Previous research has established that the Fgf/Mapk pathway exerts inhibitory effects on Smad1 via phosphorylation at the linker region which prime Smad1 to subsequent GSK3β-mediated phosphorylation leading to its polyubiquitination, and degradation (De Robertis & Kuroda, 2004, Sapkota et al., 2007). In contrast, our current and prior investigations have unveiled a novel stimulatory role of Fgf /Mapk/Xbra signaling, particularly through xbra’s involvement in Smad1 nuclear translocation and the expression of various myogenic markers, giving rise to doubt about alternate fate of Smad1 in presence of Xbra. To ascertain whether Xbra can also impede Smad1 polyubiquitination and degradation, we performed a series of experiments. Initially, we conducted co-immunoprecipitation assays involving endogenous Smad1 and flag-ubiquitin in the presence or absence of xbra. Our results, depicted in Fig. 4A, demonstrate a significant reduction in binding between endogenous Smad1 and flag-ubiquitin in the presence of Xbra. Similarly, the formation of complexes involving WT HA-Smad1 and flag-ubiquitin decreased in the presence of xbra, as illustrated in Figure 4B. Notably, flag-smad1(4SD3SD) represents the most vulnerable form for polyubiquitination and degradation(Sapkota et al., 2007). To investigate whether xbra’s interaction with flag-smad1(4SD3SD) confers protection, we conducted co-immunoprecipitation experiments involving flag-smad1(4SD3SD) and endogenous ubiquitin in the presence or absence of xbra. Our findings, shown in Fig. 4C, indicate a substantial reduction in the formation of complexes between endogenous ubiquitin and flag-smad1(4SD3SD) when xbra is present. Subsequently, we explored Smad1 degradation profile in the presence of xbra through a time-dependent Western blot (WB) assay following cycloheximide (CHX) treatment. We injected flag-ubiquitin in the presence or absence of xbra and assessed endogenous smad1 levels over time. Fig. 4D demonstrates that after 3 hours of CHX treatment, the presence of ubiquitin resulted in a marked reduction in smad1 levels compared to the control, while the injection of xbra sustained smad1 levels to a greater extent even in presence of ubiquitin. Moreover, we observed that the presence of Xbra significantly maintained the levels of flag- smad1(4SD3SD) for up to 6 hours compared to the control in CHX-treated samples (Fig. 4E). Following Fgf /Mapk-mediated linker region phosphorylation, Smad1 becomes primed for GSK3β-mediated phosphorylation, leading to polyubiquitination and degradation (Verheyen, 2007). To investigate whether Xbra could also stabilize flag-smad1(4SD3SD) in the presence of GSK3β, we conducted a time-dependent WB assay with CHX treatment in the presence or absence of Xbra. Remarkably, Xbra also maintained the levels of flag-smad1(4SD3SD) in GSK3β-injected samples (Fig. 4F), mirroring the effects of the GSK3β inhibitor LiCl (Fig. 4G). These findings underscore the intricate molecular mechanisms involving xbra, resulting in the stabilization of Fgf/Mapk/GSK3β-mediated linker region phosphorylated Smad1 through the formation of the Xbra-Smad1 complex. Furthermore, this research sheds light on the cooperative molecular interplay between Fgf /Mapk/Xbra signaling and Bmp/Smad1 output in the presence of Mapk/Erk –mediated phosphorylation at Smad1 linker region, providing deeper insights into the complex regulatory network at play.

**Fig. 4.**
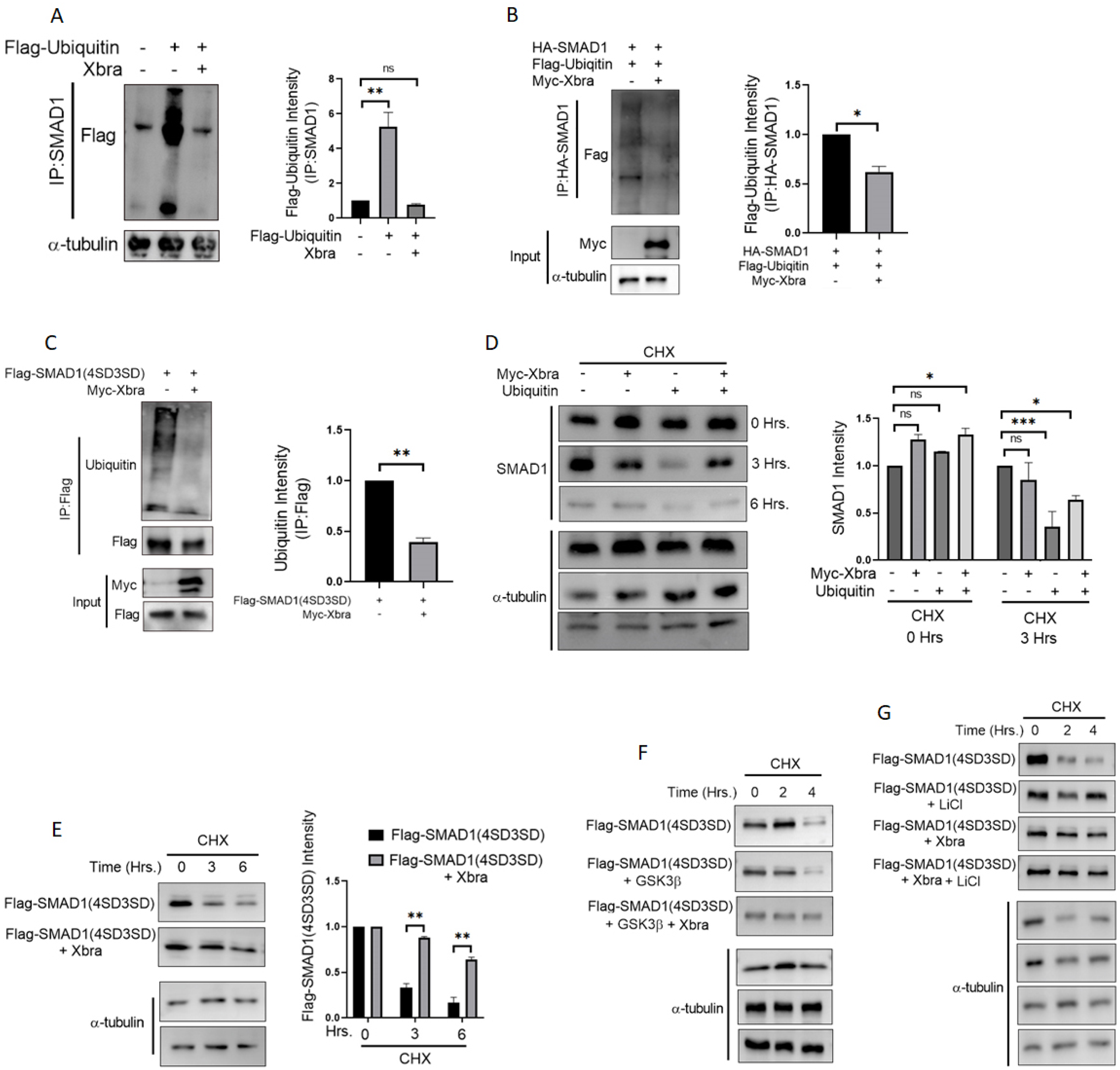
Xbra inhibits the polyubiquitination and degradation of endogenous Smad1 as well as a phospho-mimetic Smad1 Mutant. (A) Embryos injected with flag-ubiquitin with or without myc-xbra harvested at stage 10.5 for co-immunoprecipitation analysis using anti-smad1 and anti-flag antibodies. α-tubulin served as loading control. Right: Flag-Ubiquitin level were quatified by photo shop software. This experiment was repeted two times with consistemce results. (B) Embryos injected with HA-smad1(WT) and Flag-ubiquitin with or without myc-xbra harvested at stage 10.5 for co-immunoprecipitation analysis using anti-ha and anti-flag antibodies. α-tubulin served as loading control. Right: Flag-Ubiquitin level were quatified by photo shop software. This experiment was repeted two times with consistemce results (C) Embryos injected with Flag-smad1(4SD3SD) with or without myc-xbra harvested at stage 10.5 for co-immunoprecipitation analysis using anti-flag and anti-ubiquitin antibodies. Right: Endogenous Ubiquitin level were quatified by photo shop software. This experiment was repeted two times with consistemce results (D) Embryos injected with myc-xbra and ubiquitin alone or in combination were treated with 200μM/ml of cycloheximide at stage 10.5 and harvested at 0 hr. 3 hr. and 6 hr. for WB analysis using anti smad1 antibody. α-tubulin served as loading control. Right: Endogenous smad1 level were quatified by photo shop software. This experiment was repeted two times with consistemce results (E) Embryos injected with flag-smad1(4SD3SD) with or without myc-xbra were treated with 200μM/ml of cycloheximide at stage 10.5 and harvested at 0 hr. 3 hr. and 6 hr. for WB analysis using anti-flag antibody. α-tubulin served as loading control. Right: Flag-smad1 level were quatified by photo shop software. This experiment was repeted two times with consistemce results (F) Embryos injected with flag-smad1(4SD3SD), myc-xbra and myc-xbra alone or in combination were treated with 200μM/ml of cycloheximide at stage 10.5 and harvested at 0 hr. 2 hr. and 4 hr. for WB analysis using anti-flag antibody. α-tubulin served as loading control. (G) Embryos injected with flag-smad1(4SD3SD) and with or without xbra were treated with 200μM/ml of cycloheximide and Licl at stage of 10.5 and harvested at 0 hr. 2 hr. and 4 hr. for WB analysis using anti-flag antibody. α-tubulin served as loading control.

### smad4 is required for smad1/Xbra complex formation and is critical for Xbra mediated nuclear localization of phospho-mimetic smad1 mutant and lateral mesoderm marker induction

The activation of smad1 by BmpR leads to the formation of a complex with the co-Smad, Smad4, followed by translocation into the nucleus (Chen et al., 2004). To investigate smad4 role in the formation and activity of the Xbra-smad1 complex, we conducted gain and loss of function assays using smad4 morpholino (smad4(MO)) and HA-smad4. As depicted in Fig. 5B, the absence of Smad4 significantly reduced the formation of the complex between endogenous smad1 and myc-Xbra, while the injection of HA-smad4 successfully rescued the smad1-Xbra complex formation. This observation was further validated with flag-smad1(4SD3SD), where we found that the formation of the Xbra-flag-smad1(4SD3SD) complex also relies on the presence of Smad4 (Fig. 5C). Additionally, we investigated whether the nuclear localization of the Xbra-flag-smad1(4SD3SD) complex depends on the presence of Smad4. We observed a significant decrease in the nuclear localization of flag-smad1(4SD3SD) in the absence of Smad4, even though the majority of Xbra was still located in the nucleus (Fig. 5D). This might be due to the reduced formation of the Xbra-flag-smad1(4SD3SD) complex under Smad4MO conditions. Furthermore, we conducted qPCR analysis, which revealed that the presence of Smad4MO inhibited the Xbra/flag-smad1(4SD3SD)-mediated expression of myogenic markers, including *myf5*, MyoD, *myogenin*, and muscle actin (*m-actin*) and the injection of Smad4 successfully rescued the expression of these markers (Fig. 5E-H). These findings underscore the essential role of Smad4 in facilitating the formation of the Smad1-Xbra complex, its nuclear localization, and its transcriptional activity.

**Fig. 5.**
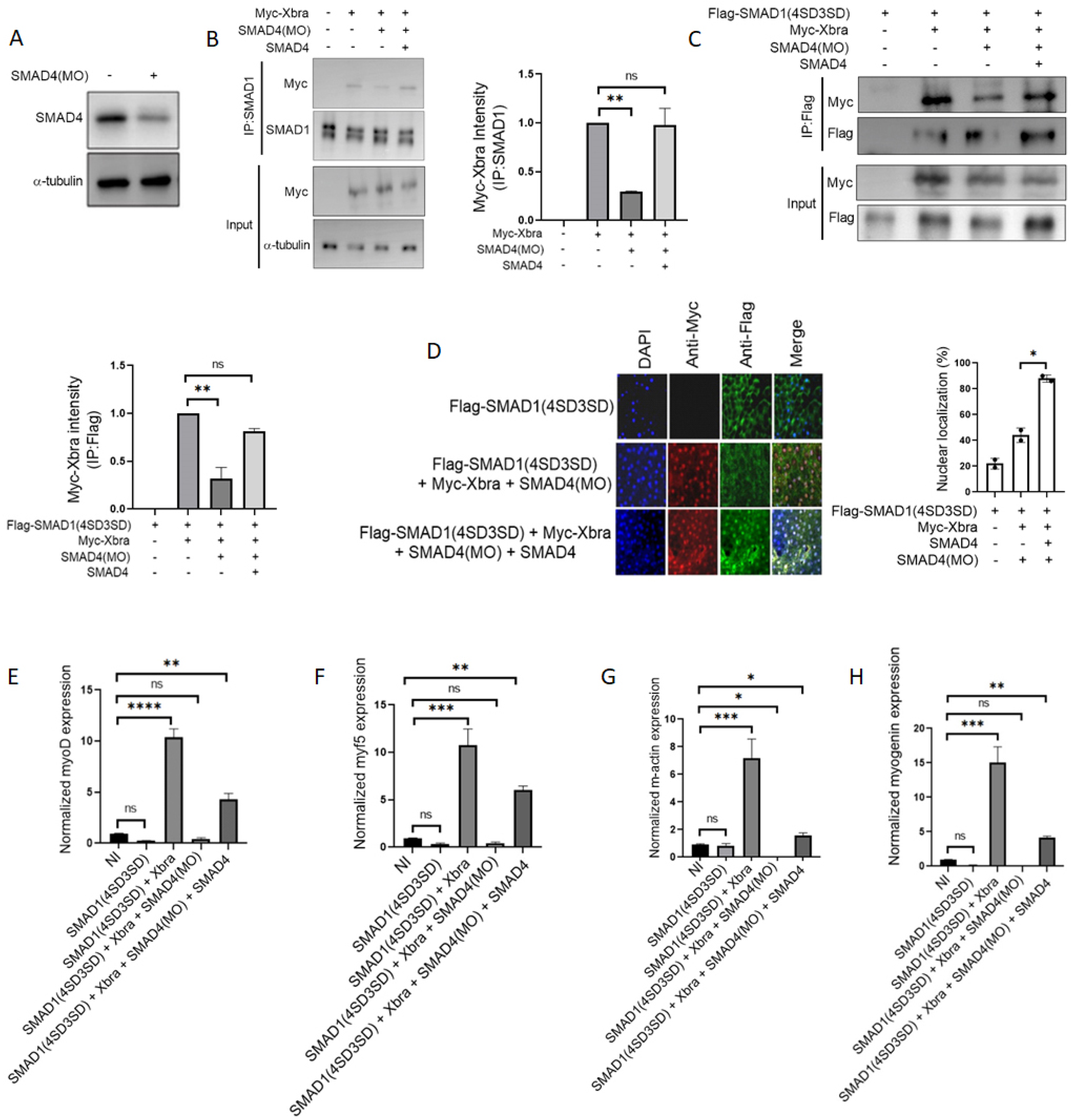
Smad4 plays a crucial role in the formation of the Smad1/Xbra complex, and it is essential for facilitating the nuclear localization and transcription activity of the phospho-mimetic Smad1 mutant mediated by Xbra. (A) Embryos injected with smad4MO harvested at stage 10.5 and subjected for WB analysis using anti-smad4 antibody. (B) Embryos injected with myc-xbra and smad4 with and without smad4MO harvested at stage 10.5 for co-immunoprecipitation analysis using anti-smad1 and anti-myc antibodies. α-tubulin served as loading control. Right: Myc-Xbra level were quatified by photo shop software. This experiment was repeted two times with consistemce results (C) Embryos injected with Flag-smad1(4SD3SD), myc-xbra and smad4 with and without smad4MO harvested at stage 10.5 for co-immunoprecipitation analysis using anti-Flag and anti-myc antibodies. Right down: Myc-Xbra level were quatified by photo shop software. This experiment was repeted two times with consistemce results (D) Subcellular localization of flag-smad1(4SD3SD) in Xenopus animal cap explant from embryos injected with flag-smad1(4SD3SD), myc-xbra and smad4 with or without smad4MO. Animal cap dissected at 9.5 stage and grown until stage 10.5, fixed and then subjected to immunofluorescence assay using primary antibody against flag-tag and myc-tag. Anti-mouse Alexa-fluor 488 and anti-rabbit Alexa fluor 594 secondary antibody (Invitrogen) was used. Blue: DAPI, Green: flag tag, Red: myc-tag. Scale bar 20X. This experiment was repeated two times with consistent results. Right: Statistic data of smad1 nuclear translocation ratio. (E-F) qRT-PCR Analysis of the gene expression levels of early markers in animal cap explants from embryos injected with smad1(4SD3SD), xbra and smad4 with or without smad4MO. qRT-PCR values were determined from the ΔΔCt for the target genes relative to odc. Significantly different results for treatments used one-way ANOVA with Graph Pad Prism; P < 0.05. (G-H) qRT-PCR Analysis of the gene expression levels of late markers in animal cap explants from embryos injected with smad1(4SD3SD), xbra and smad4 with or without smad4MO. qRT-PCR values were determined from the ΔΔCt for the target genes relative to odc. Significantly different results for treatments used one-way ANOVA with Graph Pad Prism; P < 0.05.

**Fig. 6.**
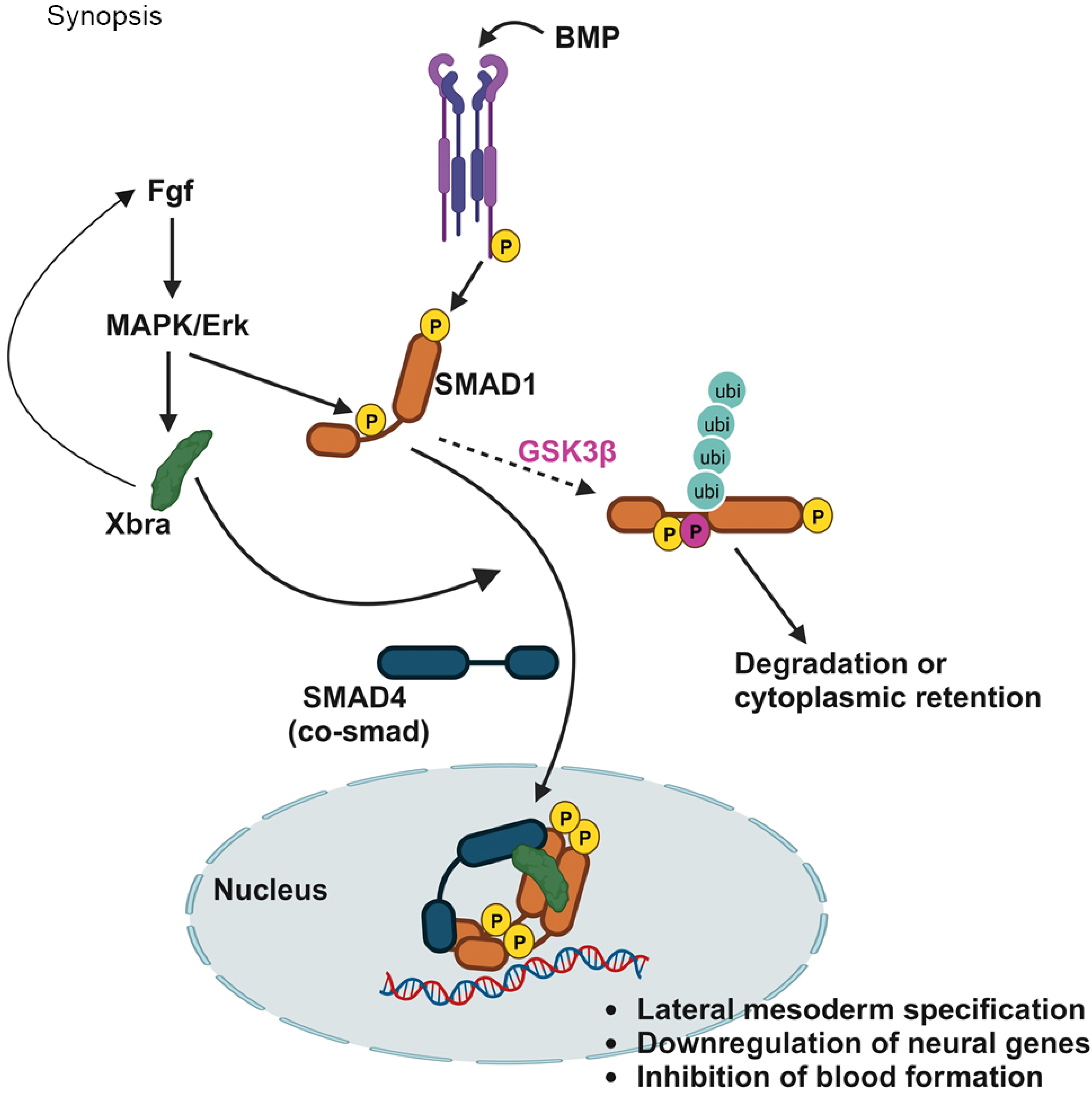
The proposed model. Followed by BMPR mediated c-terminal phosphorylation of smad1, FGF input at the linker region of smad1 further promotes GSK3β primed phosphorylation leading to its ubiquitin mediated proteasomal degradation and cytoplasmic retention. In the presence of xbra which itself generated by Fgf, linker and c-terminal phosphorylated smad1 interacts with Xbra and smad4 to form Xbra/Smad1/Smad4 trimeric complex, preventing Smad1 ubiquitin-mediated proteasomal degradation and promoting its nuclear localization and involved in upregulation of key mesodermal genes and the concurrent downregulation of neural genes and blood formation.

## Discussion

In our current investigation, we have introduced a novel regulatory pathway associated with the Bmp/Smad1 and Fgf/Mapk/Xbra signaling cascade, which may play a pivotal role in the specification of lateral mesoderm in *Xenopus* embryos. This study aimed to address two significant unresolved questions. Firstly, we sought to elucidate the dual nature of Fgf input to the Bmp pathway, whether it acts as an stimulator or inhibitor. Our hypothesis posits that this outcome depends largely on the presence of other factors. In the marginal zone of *Xenopus* embryos, the high concentration of Xbra appears to determine the fate of c-terminal and linker region phosphorylated Smad1 by forming a Xbra-Smad1 complex (Fig. 2A, B, and D). This complex, along with Smad4, escapes the Smad1 degradative pathway (Fig. 3A-G) and localizes within the nucleus (Fig. 1A and B). In contrast, the absence of Xbra in the ectodermal portion, coupled with the presence of a potent Bmp inhibitor at the dorsal-most region of the marginal zone, appears to regulate the Bmp/ Smad1 outcome differently. Secondly, we explored the significance of this cooperative regulation between Fgf/Mapk/Xbra and Bmp/ Smad1 pathways. Our findings indicate a potential role for this regulation in driving lateral mesoderm development at the expense of hematopoietic progression (Fig. 1 and 2). The varying levels of different Fgfs in different parts of the embryo during development could contribute to the differential outcome is a matter of further investigation.

The conventional Bmp gradient model has been presented to explain the dorsal-to-ventral asymmetry in mesoderm development in *Xenopus* (Dale & Jones, 1999, Dosch, Gawantka et al., 1997). According to this model, mesoderm specification occurs in two sectors within the marginal zone. One end, known as the Spemann organizer, secretes various Bmp inhibitors, including chordin, noggin, and xnr3, creating a Bmp gradient with its lowest concentration at the organizer end (dorsal) and the highest at the opposite end, known as the counter-organizer (ventral) (Dale & Jones, 1999, Harland & Gerhart, 1997, Heasman, 1997, Lane & Smith, 1999). Although Bmp gradient do found from counter-organizer to organizer end however, several studies have raised questions about the Bmp gradient model of mesoderm specification. Firstly, ventral blood island patterning occurs not only from the ventral-most part of the marginal zone but also from all meridian parts of the gastrula marginal zone (Lane & Smith, 1999, Mills, Kruep et al., 1999, Tracey, Pepling et al., 1998). Additionally, the region termed the ventral marginal zone specifies both posterior ventral (VBI) and posterior dorsal (somite) tissues in tadpole development (Kumano & Smith, 2002). Finally, the marginal zone’s tissue patterning has been observed to be polarized into vegetal and animal ends, with VBI (ventral mesoderm) originating from the vegetal end and somite (dorsal mesoderm) from the animal end of the marginal zone (Kumano, Belluzzi et al., 1999, Lane & Smith, 1999). Fgf expression has also been found to be polarized along the animal/vegetal axis in the marginal zone, and the injection of dominant-negative Fgf receptor in counter-organizer explants leads to expanded *globin* expression throughout the explant rather than being restricted to the vegetal end (Kumano & Smith, 2002). Consistent with the expression polarity of Fgf, Mapk activation is also polarized along the animal/vegetal axis of the marginal zone, with the highest expression observed toward the animal pole and very low levels in the vegetal pole (Kumano & Smith, 2000).

Fgf signaling is essential for Mapk activation, and the expression of three Fgf family members (eFgf, Fgf2, and Fgf8) has been detected predominantly in the animal polar region of the marginal zone, correlating with activated Mapk (Kumano & Smith, 2002). Similarly, several other mesoderm markers follow this animal/vegetal expression polarity in the marginal zone. MyoD is expressed throughout the marginal zone from the organizer to the counter-organizer, except in the vegetal end and Spemann organizer, and its expression depends on the presence of the Fgf pathway (Kumano & Smith, 2000). Loss of eFgf in the marginal zone results in the loss of *myoD* expression. Similarly, xbra exhibits a similar expression pattern to *myoD* in the marginal zone, extending through the Spemann organizer. The expression of *myoD* in the counter-organizer may give rise to presumptive caudal somite rather than ventral mesoderm in the animal pole end of the marginal zone(Kumano & Smith, 2000). The expression patterns of Fgf/Mapk/Xbra, along with Bmp/Smad1 in the marginal zone, suggest the possibility of their cross-talk in somite and VBI development in *Xenopus* embryos, a phenomenon that remains largely obscure.

Our present study sheds light on the integration of the Fgf/Mapk/Xbra and Bmp/ Smad1 pathways at the molecular level during somite formation in *Xenopus* embryos. Elevated expression of *myoD*, *myf5*, *myogenin*, and muscle-actin *(m-actin)* in animal cap explants overexpressed with Bmp and eFgf suggests a cooperative interplay between Fgf and Bmp in mesodermal differentiation. Furthermore, our findings indicate that the constitutive linker and c-terminal phosphorylated form of Smad1 can upregulate lateral mesodermal markers only in the presence of Xbra, suggesting the cooperative input of Fgf at the Smad1level via Xbra. Additionally, our findings illustrate the mechanistic view of Fgf/Mapk/Xbra cooperative interplay with Bmp/Smad1 via xbra in lateral mesodermal specification, suggesting Fgf/Mapk input to Smad1 promotes xbra-Smad1 interaction, protecting Smad1 from polyubiquitination and degradation and inducing Smad1 nuclear localization. This finding may indicate the requirement for a fine balance of Bmp/Smad1 activity in VBI and somite formation. In the animal pole region of the marginal zone, the inhibitory input from Mapk, along with the stimulatory input from xbra, may provide the necessary moderate Bmp/Smad1 activity for somite formation. At the vegetal pole end of the marginal zone, the absence of Fgf/Mapk activity may promote hyperactivity of Bmp/Smad1, required for VBI formation.

Our past research strongly supports that xbra plays a crucial role in the interaction between Bmp and xvent1 during mesodermal specification in Xenopus embryos. Knockdown of xvent1 promotes neurogenesis in animal cap explants treated with basic Fgf. Xbra and Smad1 cooperate in modulating the transcription of xvent1, with Xbra knockdown leading to increased neuroectoderm formation (Kumar et al., 2018, Lee, Lee et al., 2011, Yoon et al., 2014). This study identifies Xbra as a key determinant of Bmp/ Smad1 pathway during mesodermal specification. We explored the molecular pathway involving Fgf/Xbra and Bmp/Smad1, highlighting the critical role of Xbra–Smad1 complex formation. Fgf-mediated phosphorylation of Smad1 enhances its affinity for Xbra binding and further preventing its degradation and promoting stability (Fig. 4) by preventing polyubiquitination and degradation. Xbra also facilitates Smad1 nuclear localization and transcriptional activity in the presence of smad4. Our findings provide insights into the molecular mechanisms of Fgf/Mapk/Xbra collaboration, indicating a regulatory role of Fgf in the Bmp pathway during mesodermal specification in Xenopus embryos, in contrast to its inhibitory role in neural patterning. These interactions may contribute to diverse aspects of embryonic development.

At last, our study provides insights into the cooperative crosstalk between Fgf/Mapk/Xbra and Bmp/Smad1 signaling pathways during mesodermal specification in *Xenopus* embryos. Contrary to the usual belief that Fgf only inhibits Bmp, this study reveals a more complex role for Fgf in connection with Bmp4/Smad1, which merits further evaluation

## Material and method

### Embryo Manipulation

This study was conducted considering all ethical regulation according to the guidelines of Institutes of Laboratory Animal Resources, Hallym University that specifically work for laboratory animal maintenance. Eggs were obtained from female *Xenopus* laevis by injecting 700 U of human chorionic gonadotropin (HCG). Eggs were in-vitro fertilized using macerated testis. Fertilized embryos were dejelled in 2% L-cysteine (Bio Basic) solution pH 7.8. Embryos were cultured in 0.33% modified Ringer solution. Stage determination of embryo were done according to Nieuwkoop and Faber’s table of development. Embryos were microinjected at one cell stage in the animal pole region. For animal cap explant, explants were dissected at 8-8.5 stage and cultured until stage 10.5 or stage 24 in L-15 media (gibco) containing bovine serum albumin (BSA, BOVOGEN) and 50 μg/ml of gentamicin then explants were subjected for further analysis.

### Plasmids

The Flag-smad1 WT construct was produced by inserting inserting the coding region of human Smad1 into the pSP64T expression vector. The mutant Flag-smad1(4SD3SD) construct was achieved by using Flag-smad1 WT. The amino acid changes from S to D was achieved by using by site-directed mutagenesis using Pfu DNA polymerase (Elpis Biotech) and the primers, 5’-CCTCACGACCCCAATAGCAGTTACCCAAACGACCCT-3’ (hSmad1-3SD-Forward), 5’-AGGGTCGTTTGGGTAACTGCTATTGGGGTCGTGAGG-3’ (hSmad1-3SD-Reverse), 5’-CACGACCCCACCAGCTCAGACCCAGGAGACCCTTT -3’ (hSmad1-4SD-Forward) and 5’-GGGTCTCCTGGGTCTGAGCTGGTGGGGTCGTGAG-3’ (hSmad1-4SD-Reverse). Myc-xbra was cloned into the pCS2+ expression vector (Lee et al., 2011)

### Morpholino oligos

The smad4 anti-sense morpholino oligos (MOs) were purchased from Gene Tools. The sequences of Smad4 MOs are as follows:

XSmad4-α-MOs: 5’-TGTTTGTGATGGACATATTGTCGGT-3’

XSmad4-β-MOs: 5’-GGGTCAGAGACATGGCCGGGATCTC-3’

Both α and β smad4 MO were mixed in equal amount and injected for knock down.

### mRNA synthesis

Capped synthetic mRNAs were produced by using Ambion mMessage mMachine SP6 (AM1340) kits (Thermo Scientific, Inc.). Flag-smad1 (WT), Flag-smad1(4SD3SD), Flag-smad1(4SA3SD), HA-Smad4 and Bmp4 were linearized with ASP718 and their respective mRNA was synthesized with SP6 RNA polymerase. Myc-Xbra was linearized with sal1 and their respective mRNA was synthesized with SP6 RNA polymerase. eFgf was linearized with Not1 and their respective mRNA was synthesized with SP6 RNA polymerase.

### RNA isolation and quantitative RT-PCR (qPCR)

Total RNA was isolated from animal cap explants using TRIzol Reagent (Life Technologies) and treated with RNase-free DNase Ι (Roche) to remove genomic DNA contamination. Approximately 4 μg of RNA was used for reverse transcription by using oligo DT and SuperScript IV reverse transcriptase (Invitrogen). qPCR was performed in a standard 20 μl mixure with KAPA SYBER FAST qPCR Master Mix using an Applied Biosystems StepOnePlus Real-Time PCR System. All the real-time values were averaged and compared using the threshold cycle (CT) method, in which the amount of target RNA (2 − ΔΔCT) was normalized against the endogenous expression of ODC (ornithine decarboxylase) (ΔCT) All qPCR reactions were repeated three time using independent samples to present data with standard deviations and statistical significance.

### Immunoprecipitation and Western blot analysis

For co-immunoprecipitations and Western blotting of endogenous and ectopically expressed proteins in *Xenopus* embryos were harvested at the stage 10.5. Embryos were homogenized in lysis buffer (50 mM Tris (pH 7.6), 1% NP-40, 0.5% Sodium deoxycholate, 1% Triton X-100, 150 mM NaCl, 5 mM EDTA, 0.1% SDS 1 mM sodium vanadate, 10 mM NaF, 1 mM PMSF, 1X Halt Protease inhibitor cocktail (Thermo Scientific REF 78430) after homogenized, lysis buffer incubated in respective antibodies for overnight and further incubated in agarose beads. The beads were washed and lysate were boiled with loading buffer. Proteins were separated via sodium dodecyl-sulfate polyacrylamide gel electrophoresis and transferred to membranes. These were blotted with proper antibodies and visualized with ECL.

### Immunofluorescence

The injected and un-injected embryos were dissected at stage 9.5 for immunofluorescence assay. Dissected ACs were fixed in 4% paraformaldehyde for 2 h and incubated in 2% BSA in phosphate-buffered saline (PBS) containing 0.5% Triton-X for 1 h. The ACs were incubated with primary antibodies overnight at 4 °C, washed with PBS, and incubated with fluorescent secondary antibodies for 2 h at room temperature. The ACs were then mounted, and images were obtained and analyzed using confocal microscopy (CLSM II, Carl Zeiss LSM-710) (Carl Zeiss, Oberkochen, Germany).

### Statistical analysis

Statistical comparisons were done by using ordinary one-way analysis of variance (ANOVA) analysis followed by multiple comparisons or Student’s t test by using GraphPad Prism. P < 0.05 was considered to indicate a statistically significant difference.

## Acknowledgements

This study was supported by the Basic Science Research Program of the National Research Foundation of Korea (NRF), funded by the Ministry of Education, Science, And Technology of Korea (2018M3C7A1056285, 2021R1A4A1027355, and 2021M3H9A1097557).

